# Modeling population control via tunable sex ratio distorter gene drives in *Aedes aegypti*, an arboviral vector with homomorphic sex chromosomes

**DOI:** 10.64898/2026.07.05.736587

**Authors:** Lauren M. Childs, Sara Shabani, Uwe C. Täuber, Zhijian Tu

## Abstract

*Aedes aegypti*, a major vector of arboviruses, belong to subfamily Culicinae, a diverse group of mosquitoes with homomorphic sex-determining chromosomes. Males are the heterogametic sex with a dominant male-determining locus (M locus). The M locus and its counterpart m locus are embedded in a region of suppressed recombination, with a large portion of this recombination desert showing significant molecular differentiation despite homomorphy. We developed a mathematical framework to examine M-linked genome editors that specifically target the m-chromosome during spermatogenesis, mimicking the naturally occurring sex ratio distorters (SRDs) in Culicinae that produce male-biased meiotic drives. Unlike previous models for species with heteromorphic sex chromosomes (e.g., X and Y), we incorporate features stemming from the homomorphic nature of the *Ae. aegypti* sex chromosomes such as varied linkage to the M locus, making the degree of super-Mendelian inheritance readily tunable. We evaluated *in silico* SRDs with a range of M-linkage and editing efficiencies and established the theoretical foundation for developing highly efficient SRDs that outperform several methods of population suppression. These SRDs can be tuned to mitigate impact on a neighboring population. The framework developed here is suitable for exploring SRD-mediated genetic biocontrol of numerous pest species with homomorphic sex chromosomes.

## 1 Introduction

Mosquitoes comprise the deadliest animal in the world, in particular, *Anopheles gambiae*, a major vector of malaria, and *Aedes aegypti*, which transmits dengue, chikungunya, and Zika viruses [1]. Frontline prevention of disease outbreaks from these pathogens depends mainly on effective vector control. Historically this was dominated by killing of the vector through the application of insecticide, the effectiveness of which is hindered by recent increases in insecticide-resistance. Insecticide usage also has the potential to kill or harm non-target species in the environment. Thus, species-specific genetic biocontrol methods are being developed with the goal to either suppress the target mosquito population or to modify the population so the mosquitoes harbor more favorable traits, such as resistance to the pathogen they are known to transmit (reviewed in [2]). While population modification relies on gene drives to bias the inheritance of an effector gene(s) or allele(s) to achieve its fixation in the target vector population, the most common strategies for population suppression involve the release of the non-biting males that are made sterile through irradiation (as in classic Sterile Insect Technique or SIT), genetic manipulation (e.g., precision guided SIT or pgSIT), or Wolbachia infection (as in the Incompatibility Insect Technique or IIT) (reviewed in [2, 3, 4]).

Sex ratio distortion, reducing the number of female offspring relative to male offspring, is a potentially more powerful alternative method to achieve population suppression. Reducing the number of females has an outsized benefit as females are the main contributors to the population size and only female mosquitoes can transmit pathogens. Genetically modified *An. gambiae* lines were generated through the insertion in the autosome of an engineered endonuclease or guided CRISPR/Cas9 that targets the rDNA cluster on the X-chromosome for selective cleavage of the X-chromosome during spermatogenesis. These ‘X-shredding’ lines produced predominantly Y-bearing sperm, resulting in as much as 97.4% male biased progeny [5, 6]. These synthetic X-shredders are sex ratio distorters (SRDs), and they significantly more efficient at population suppression than sterile males due to the multi-generational effect [7, 8]. As the above-mentioned X-shredders are located on an autosome, their inheritance follows the Mendelian ratio. However, when the X-shredder is inserted onto the Y chromosome, it becomes a gene drive as its inheritance is *>*50% due to the destruction of the X-bearing sperm. Thus, Y-linked X-shredders and other Y-linked male-biased SRDs are much more powerful at population suppression compared to their autosomal counterpart according to mathematical models [9, 10, 11, 8], with some requiring only a single release of less than 1% of the target population, and are thus potentially invasive [12]. However, Y-linked X-shredders have not been reported, likely due to meiotic sex chromosome inactivation which will result in silencing of the Y-linked shredder during meiosis in organisms such as *An. gambiae* [13, 14].

Unlike the well differentiated X and Y chromosomes in *Anopheles* mosquitoes, the sex chromosomes of *Ae. aegypti* consist of a pair of homomorphic sex chromosomes (chromosome 1) that are structurally indistinguishable from autosomal chromosomes except around the sex-determining locus (reviewed in [15]). In *Ae. aegypti*, maleness is determined by a dominant male-determining factor (M factor) *Nix* [16] found within a small male-determining locus (M locus). The m locus, the counterpart of the M locus, is characterized by the lack of *Nix* and other M locus genes (Figure 1; reviewed in [15]). The sex locus is in an *≈*100 Mb region of suppressed recombination, and a significant portion of this recombination desert shows significant molecular differentiation between the M-bearing and m-bearing chromosomes despite similarity in form, i.e. homomorphy [17, 18, 19, 20, 21]. Some strains of *Ae. aegypti* showed naturally occurring sex ratio distortion, producing a higher percentage of males than females in the offspring [22, 23]. The increased male percentage may result from m-chromosome breakage mainly at one of four sites during male meiosis [24]. The locus associated with sex ratio distortion was mapped close to the M locus [25], although efforts to identify the distorter have not been successful (e.g., [26]).

**Figure 1.**
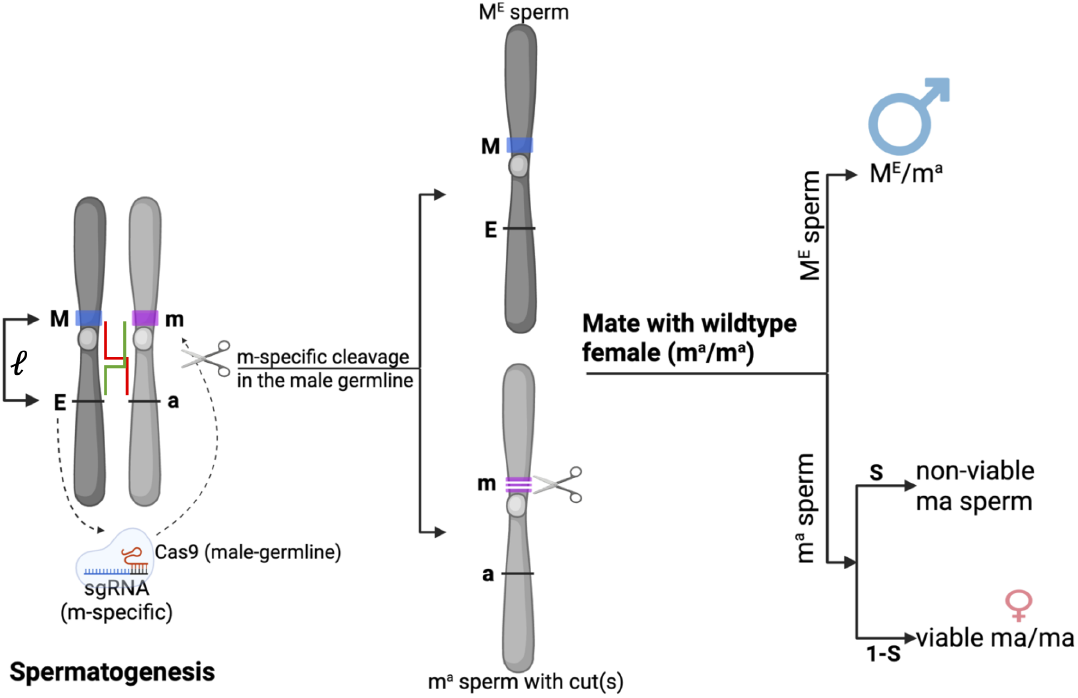
A tunable sex ratio distorter in *Aedes aegypti*. During spermatogenesis (left), targeted sex chromosome cleavage (middle) results in male-biased progeny (right). M, the male-determining locus which contains a dominant male-determining factor (M factor) *Nix* [16] that initiates male development in *Ae. aegypti* ; m, the counterpart of the M locus characterized by the lack of *Nix* and other M locus genes; E, a synthetic sex ratio distorter that expresses Cas9 in the male germline and single guide RNAs (sgRNAs) that specifically target the m locus; a, a wildtype allele of E; *ℓ*, the genetic linkage between M and E with red and green lines indicating possible recombination; *s*, the rate at which the m chromosome is “shredded” during spermatogenesis resulting in a non-viable m sperm unable to fertilize an egg (pre-zygotic lethality). *s* is analogous to the rate of X-shredding as described in *Anopheles gambiae* and *Drosophila melanogaster* (e.g., [12]). We explore the effects of continuous M-E linkage, denoted by M^E^, (50-100% with *ℓ ∈* [0.5, 1]) and relative effectiveness of shredding (*s ∈* [0, 1]) as novel features to develop tunable sex ratio distorters for efficient and confined genetic biocontrol of *Ae. aegypti*. Only non-recombinant gametes are shown.

*Ae. aegypti* is but one of the approximately 3200 mosquitoes in the Subfamily *Culicinae* that have homomorphic sex chromosomes (e.g., [27]). Sex chromosome homomorphy is also common in other insects, fishes, amphibians, reptiles, and plants [28]. As meiotic sex chromosome inactivation results from unsynapsed chromatin [29], it is generally not expected for homomorphic sex chromosomes, potentially removing a major bottleneck in developing synthetic Y-linked (or M-linked) SRDs. Partly due to its important disease vector status, *Ae. aegypti* represents one of the best characterized homomorphic sex chromosomes (reviewed in [15] and see Section 2.1). Thus, we focused on *Ae. aegypti* in this study. We developed a mathematical framework to examine M-linked SRDs (or m-shredders) by incorporating new features stemming from the homomorphic nature of the *Ae. aegypti* sex chromosomes such as the ability to establish SRDs with varied degrees of linkage to the M locus. Using this modeling framework, we established and evaluated *in silico* SRDs with a range of efficiencies and confinement properties, which expands the toolbox to control mosquito-borne infectious diseases. We demonstrated that m-shredders of tight M-linkage (*>*98%) are similar in their effectiveness for population suppression as an m-shredder with 100% M linkage, which negates the need to insert the m-shredder into the highly repetitive M locus, thus, mitigating a significant technical challenge. The addition of this new parameter, which is relatively straight-forward to manipulate by adjusting the m-shredder insertion sites, provides potentially effective methods to modulate or balance the efficiency and invasiveness of the M-linked SRDs, which can help accommodate the varied needs facing global mosquito-borne infectious disease control programs. The framework developed in this study can be applied to numerous other species that have homomorphic sex chromosomes [28].

## 2 Methods

We develop a population-level mathematical model for the dynamics of *Aedes aegypti* mosquito populations with distinct genotypes, where each genotype is represented by two loci: one representing sex determination (M or m) and the other representing an ‘editor’ gene (E or a, where a is wildtype). For an organism to be female, both sex-determining alleles must be m, while the combination of an M and m indicates a male. The E allele in the editor gene locus represents the entire transgenic unit that expresses both the Cas9 endonuclease and the specific single guide RNAs (sgRNAs). When E consists of male-germline-specific Cas9 and sgRNA(s) that specifically targets the m locus or m-chromosome (as seen in Figure 1), it can shred the m-chromosome during spermatogenesis and functions as a SRD that confers male bias. When E is linked to the M locus (M^E^), its inheritance is non-Mendelian (Figure 1). Eight genotypes (M^E^/m^E^, M^E^/m^a^, M^a^/m^E^, M^a^/m^a^, m^E^/m^E^, m^E^/m^a^, m^a^/m^E^, and m^a^/m^a^) can result from all possible recombinant and non-recombinant gamete combinations, where the superscript represents alleles inherited together, and thus potentially linked.

### 2.1 Biological rationale for the model of M-linked m-shredders

We model M-linked m-shredders (M-linked E, represented as M^E^) as tunable SRDs (Figure 1) to achieve population suppression of *Ae. aegypti*. The feasibility of establishing these M-linked SRDs is supported by the following biological observations. First, the presence of naturally occurring M-linked SRDs in both Aedes and Culex mosquitoes (e.g., [22, 23, 30]) supports the feasibility of developing M-linked SRDs. Second, despite the homomorphic nature of the *Ae. aegypti* sex chromosomes, the *∼*1.3 Mb long M and m locus do not appear to share conserved sequences [15]. Furthermore, substantial differentiation has been characterized between the M-bearing and m-bearing chromosomes, especially in a region of suppressed recombination (at least dozens of megabases) flanking the M/m loci (e.g., [18, 19, 20, 21]). Therefore, m-specific cleavage guided by m-specific sgRNAs is feasible (Figure 1), and the naturally occurring SRD has been shown to result from m-chromosome breakage mainly at one of four sites during male meiosis [24]. Third, the homomorphic nature of the sex chromosomes in *Ae. aegypti* (e.g., [18, 20]; see Figure 1) and other *Culicinae* mosquitoes ([27]) suggests that meiotic sex chromosome inactivation is unlikely ([31]) in the subfamily *Culicinae*, removing a potential bottleneck in establishing M-linked SRDs. Fourth, male-germline-specific promoters have been identified and shown to successfully drive Cas9 expression in *Ae. aegypti* (e.g., [32, 33]). Finally, the homomorphic sex chromosomes provide the opportunity to establish SRDs with varied degrees of linkage to the M locus ([21, 34]). As shown in Figure 1, we explore the effects of continued M^E^ (M-SRD) linkage (50%-100%, *ℓ* ∈ [0.5, 1]) and relative effectiveness of shredding (*s*) as novel features to develop tunable SRDs to help efficient and confined genetic biocontrol of *Ae. aegypti*. While our model focuses on *Ae. aegypti*, we expect the framework is applicable to a wide range of other species with homomorphic sex chromosomes such as many mosquitoes in the subfamily *Culicinae* (e.g., [27]) and other organisms with homomorphic sex chromosomes including fishes, amphibians, reptiles, other insects and plants [28].

## 2.2 Model formulation

In our conceptual model framework (see Figure A1), we track the population abundance of juveniles and adults of each genotype with a fundamental time scale of a single juvenile generation. See Section A for full details on the methodology including equations and Table A1 for standard parameter values.

The abundance of offspring genotypes is determined by offspring probabilities from the individual mating pairs in combination with the abundance of the parental genotypes. Our model assumes that the editor allele targets and shreds the chromosome carrying the m alleles during spermatogensis, increasing the probability of inheriting the M allele and thus biasing the offspring toward an enhanced frequency of males. To model this, we implement a shredding efficiency parameter (*s*) that creates a bias toward male offspring. The linkage (*ℓ*) between the male-determining locus and the editor gene locus, allowed to be variable, determines the likelihood of offspring with the same combination of the sex-determining and editor gene loci as in the paternal genes.

As an extension to our model, we consider bidirectional spillover between a target population (population 1), where mosquitoes containing the editor allele are introduced, and a neighboring population (population 2) composed of a stable wildtype population. We assume that spillover between populations occurs at a constant rate proportional to the population abundance of each particular genotype. We vary the level of spillover (*σ*), but always assume it is small (*σ ≤* 5%).

### 2.3 Simulation protocol

We begin our simulations with the assumption that the mosquito population is in equilibrium with wildtype genotypes (M^a^/m^a^ and m^a^/m^a^). The wildtype population equilibrium is a balance between the birth rate and density dependence mortality experienced during the juvenile stage (with intrinsic growth rate *R*_*m*_ = 6 as discussed in Section A.2). To understand the ability of an introduced editor gene to elicit population suppression, we release male adult mosquitoes with the editor gene linked to the male-determining locus (we focus on M^E^/m^a^ release) and track how the population size and proportion of editor gene presence is altered after thirty generations unless otherwise specified. We primarily consider a single release when the population is at wildtype equilibrium, but also consider multiple releases (one per generation) of the same fraction (fixed relative to the wildtype equilibrium). All deterministic simulations are run in Matlab (R2025b) and stochastic simulations in Python. Code will be made publicly available in GitHub upon acceptance. Code for review is in a zipped supplemental file.

## 3 Results

### Population suppression possible for a wide range of M-linkage (*ℓ*) and shredding efficiency (*s*)

Our mathematical framework incorporates the impact of varied degrees of shredding and M-linkage, a feature that is relevant to species with homomorphic sex chromosomes. We first discuss pertinent results from numerically solving the deterministic system in Equation (2), but also assess certain scenarios through numerical simulation of our stochastic representation. With an autosomal m-shredder (Fig. 2, linkage of 50% or *ℓ* = 0.5, left panels), partial population suppression is readily observable with repeated release at a 0.1*×* release fraction (10% of the wildtype equilibrium male population), with the level of reduction related to the extent of shredding efficiency. With the introduction of an M-linked m-shredder, population suppression can be achieved within 30 generations for a perfect m-shredder (*s* = 1) after a single release (Fig. 2, top right panel). The results (time to elimination and final population size) are very similar for shredders with 98-100% M-linkage (*ℓ ≈* [0.98, 1]). For moderate linkage (*ℓ* = 0.8), more than 50% population suppression is seen with efficient shredding (*s >* 0.95) from a single release (Fig. 2, top second from the left panel).

**Figure 2.**
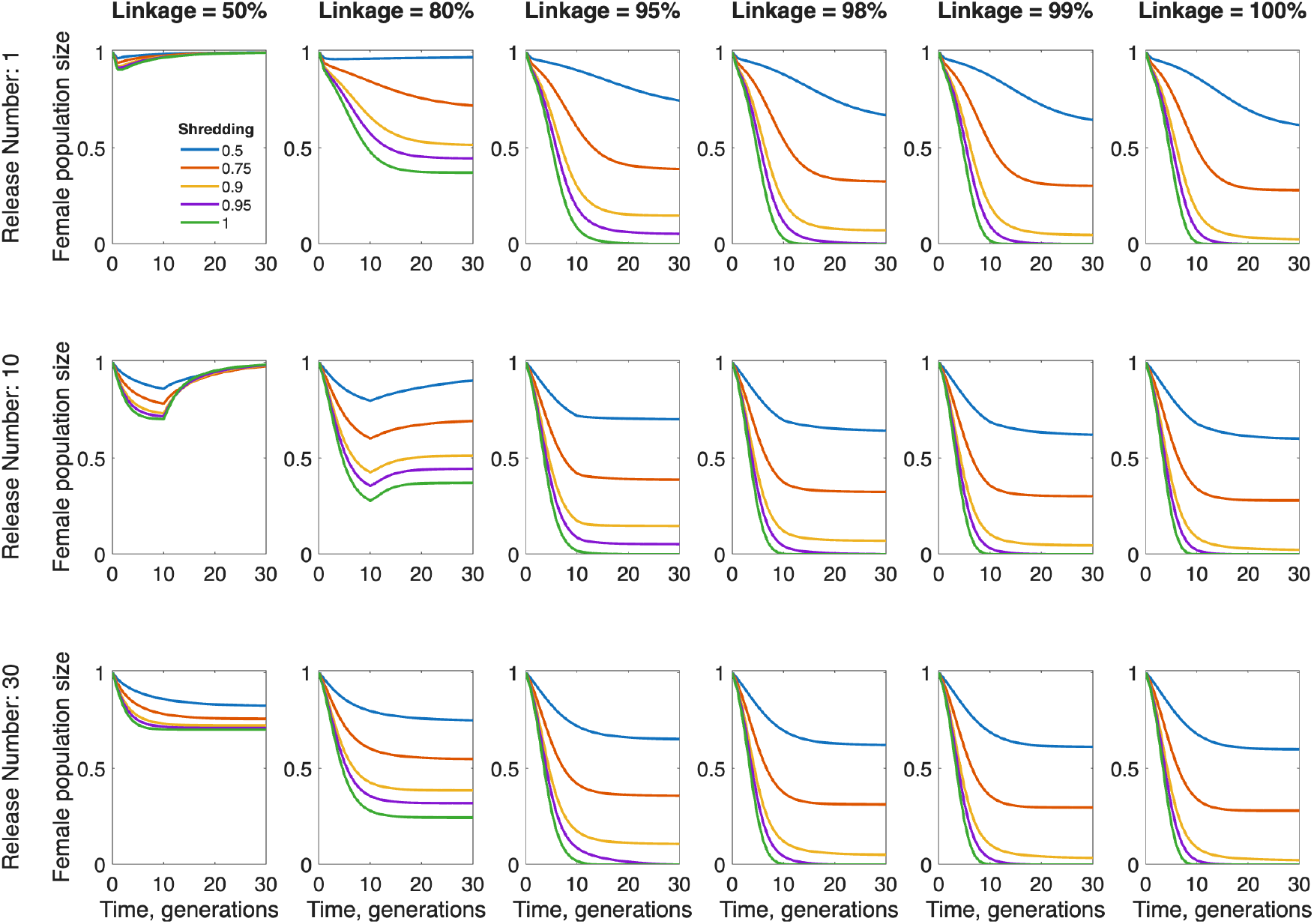
Dynamics of mosquito population across 30 generations with a single release (top panels) or repeated release (middle panels with 10 releases and bottom panels with 30 releases) of M^E^/m^a^ males under varying linkage and shredding efficiency. Relative female mosquito population size compared to a wildtype population at equilibrium following the introduction of M^E^/m^a^ males released at 0.1*×* fraction of the equilibrium male population. Linkage varies across panels from left to right: 50% (or autosomal), 80%, 95%, 98%, 99%, and 100% (fully linked). Line color denotes shredding efficiency (blue = 50%, red 75%, orange 90%, purple 95%, and green 100%). Note that m-shredders of 98-100% M-linkage provide similar levels of population suppression. Unless otherwise specified, baseline parameter values from Table A1 are used.

To assess a wider range of parameters, in Figure 3, we show the resulting female population size (relative to wildtype equilibrium) after 30 generations. Nearly complete population suppression is possible following a single release at a fraction of the wildtype male population (0.1*×*, 0.5*×*, 1*×*), when linkage and shredding efficiency are high. A slightly lower minimum linkage and shredding efficiency is needed to achieve suppression when the release fraction is larger (increasing blue area in panels left to right) and with more repeated releases (increasing blue area in panels top to bottom).

**Figure 3.**
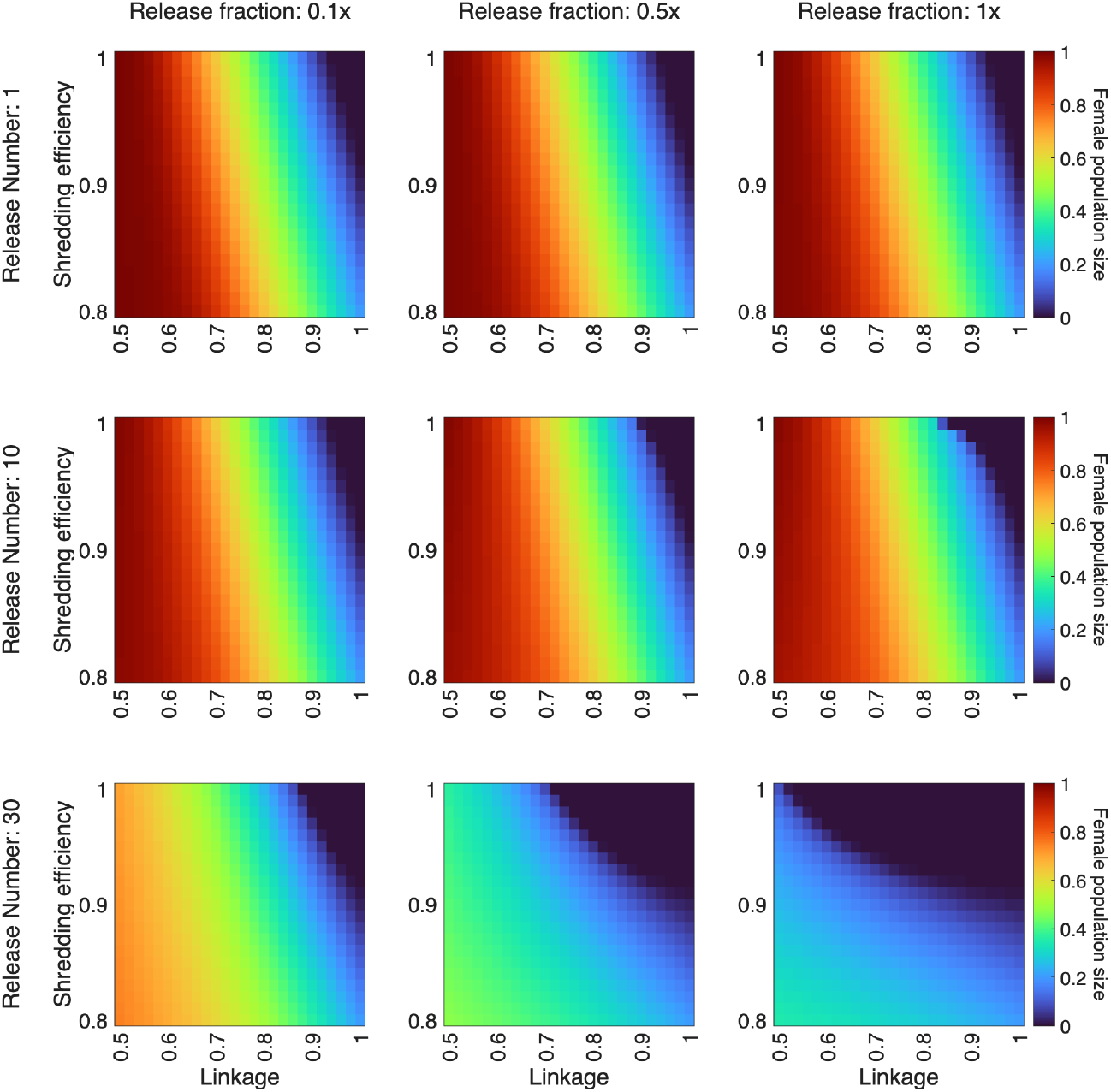
Female population (relative to wildtype equilibrium) 30 generations following a single release (top panels) or multiple repeated releases (middle panels with 10 releases and bottom panels with 30 releases) of M^E^/m^a^ males for varying linkage and shredding efficiency. From left to right, the release fraction increases (*left to right* : 0.1*×*, 0.5*×*, 1*×*). Unless otherwise specified, baseline parameter values from Table A1 are used.

With repeated release, for the same combination of linkage and shredding efficiency, smaller release fractions achieve population suppression, the extent of which is determined primarily by linkage.

These results inform the design and implementation of the M-linked m-shredders in three important ways. First, it is critical to develop effective m-shredding editors to achieve efficient population suppression. Second, the fact that m-shredders of 98% M-linkage behave similarly to those of 100% linkage greatly enhances the feasibility for highly efficient population suppression as the shredder can be inserted in a *>* 100 Mb genomic region with *≥* 98% M-linkage [21], not limited to the repeat rich M locus. Third, it is possible to modulate the effectiveness of the SRD by changing its insertion site, and, thus, its M-linkage.

### M-linked editor outperforms other population suppression techniques

Previous studies compared the effectiveness of various population suppression methods by estimating the necessary release fraction needed for repeated releases [10, 35]. We find the necessary release fraction for the M-linked m-shredder to reduce the population by 95% is much below other population suppression techniques (e.g., those found in [10, 35] ) such as Y-Linked Editor (YLE), Release of Insects carrying Dominant autosomal Lethal genes (RIDL), or autosomal X-shredder, as shown in Table 1. Optimal M-linked m-shredders with complete linkage (*ℓ* = 1) and shredding efficiency (*s* = 1) are extremely effective at population suppression even for very low release fractions. Moderate reductions in shredding and linkage (*s* = 0.92, *ℓ* = 0.99) still produce an M-linked m-shredder that outperforms optimal Y-linked Editor, optimal autosomal X-shredder and optimal RIDL. An even weaker M-linked m-shredder (*s* = 0.9, *ℓ* = 0.98) shows comparable release sizes needed for reductions to 95% suppression; however, they are unable to achieve 99% suppression due to the intrinsic population expansion by the females adults that escaped shredding.

**Table 1:** Minimum release fraction to achieve 95% or 99% suppression with 30 generations of repeated release. Supp. = suppression. ^*†*^Weak M-linked m-shredder has 90% shredding efficiency (*s* = 0.9) and 98% linkage (*ℓ* = 0.98). ^*‡*^Mid M-linked m-shredder has 92% shredding efficiency (*s* = 0.92) and 99% linkage (*ℓ* = 0.99). ^*ℓ*^Optimal M-linked m-shredder has full shredding efficiency (*s* = 1) and complete linkage (*ℓ* = 1). ^*σ*^Release fractions greater than 10*×* or less than 0.1*×* equilibrium adult male population were not tested. ^*≈*^Models used for comparison to M-linked m-shredder from [10] with identical parameterization; comparably calculated values in [10] (and [35]) are slightly lower as they are reported in [10, 35] after 36 generations of repeated release rather than 30 generations. YLE = Y-linked editor; RIDL = Release of Insects carrying Dominant autosomal Lethal genes. bi-RIDL-b is equivalent to sterile insect technique.

| Supp.<br>Level | M-linked m-shredders |  |  | Optimal<br>YLE-a* | Optimal<br>autosomal<br>X-shredder* | Optimal<br>bi-RIDL-a* | Optimal<br>fs-RIDL-a* | Optimal<br>bi-RIDL-b* |
| --- | --- | --- | --- | --- | --- | --- | --- | --- |
|  | weak<sup>†</sup> | mid<sup>‡</sup> | optimal<sup>§</sup> |  |  |  |  |  |
| 95 % | 0.100 | < 0.001<sup>#</sup> | < 0.001<sup>#</sup> | 0.046 | 0.406 | 0.446 | 0.684 | 1.317 |
| 99 % | > 10<sup>#</sup> | 0.024 | < 0.001<sup>#</sup> | 0.052 | 0.416 | 0.448 | 0.699 | 1.322 |

### Demographic stochasticity enhances the likelihood of suppression for a wide range of M-linkage and shredding efficiency

Based upon our conceptual framework for modeling m-shredder with varied degrees of M-linkage, we developed a stochastic representation which allowed us to quantify variability in population suppression, and time to local elimination, which is of particular interest and importance when population sizes are small. The combination of high linkage and high shredding efficiency were associated with the ability of the shredding allele to drive the target population to suppression with high reliability (Figs. 4 and A5, upper right corner of panels), with only slight benefits from higher release fraction. In contrast to the deterministic representation, the probability of suppression, determined by stochastic simulation, showed that intermediate linkage and shredding efficiency could still lead to population suppression after a single release (Fig. 4, bottom panels). Furthermore, the average female population size decreased and likelihood of suppression increased following repeated release for five generations (Fig. A5).

**Figure 4.**
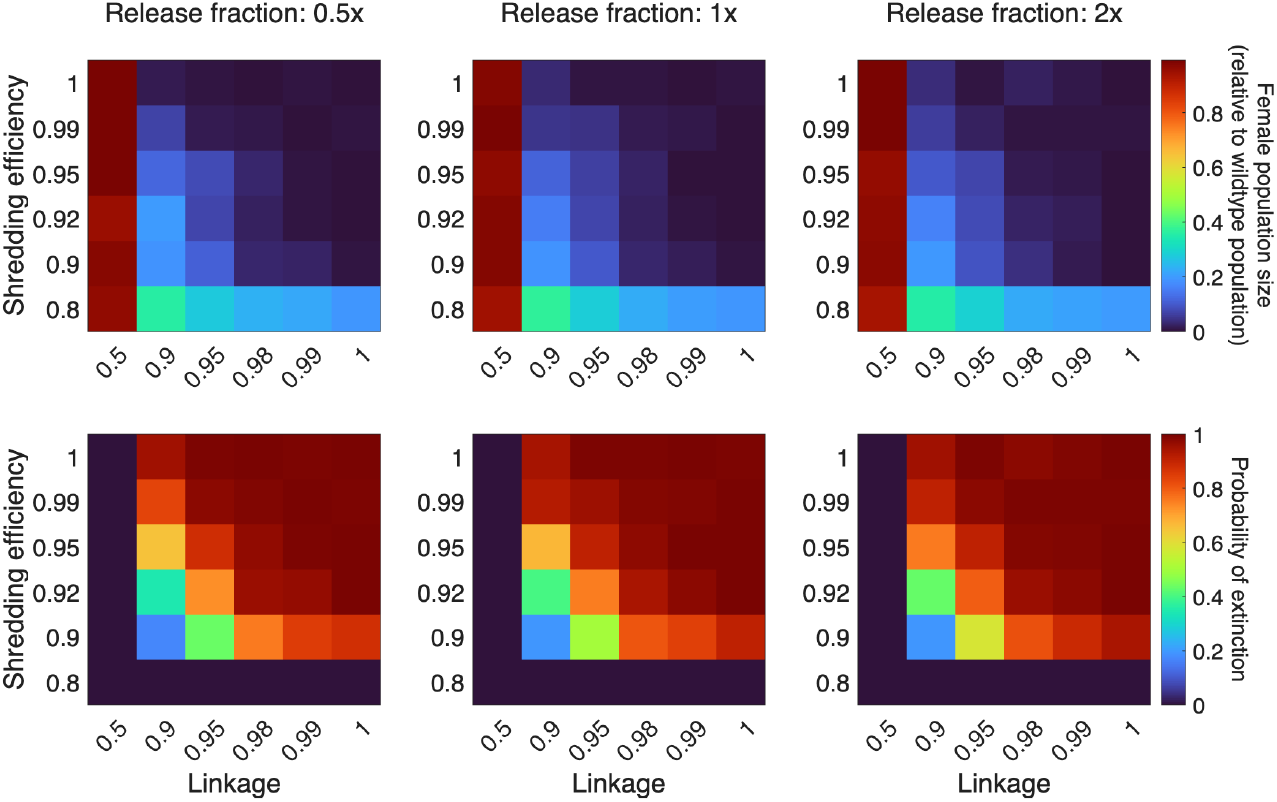
Female population size relative to wildtype equilibrium at 30 generations (top panels) and probability of suppression within 30 generations (bottom panels) following a single release of M^E^/m^a^ males for varying linkage and shredding efficiency as obtained from our stochastic representation. From left to right, the release fraction increases (*left to right* : 0.5*×*, 1*×*, 2*×*). Note the non-linear scaling of the axes. Output averaged over 200 replicate simulations with release starting after the wildtype population reached quasi-equilibrium. As fitness costs of genotypes containing the shredding allele had minimal effect on the results for the deterministic representation for 80% to 100% fitness (Fig. A6), we assumed no fitness cost for the stochastic simulations (*f*_*i*_ = 1 for all *i*). Unless otherwise specified, baseline parameter values from Table A1 are used.

### Highly robust m-shredders lead to suppression of neighboring populations

We extended our original single-population deterministic model to include a neighboring population such that we considered bidirectional spillover between the target population (population 1) and a neigh-boring population (population 2). We assumed that spillover of each genotype occurs relative to the population size of that genotype in the current population. Suppression (or local elimination) of both target and neighboring populations occurred when the M-linked m-shredder acts a strong drive, often a product of high fitness of male genotypes containing the shredding allele. This could occur even after just a single release of a highly fit genotype with strong shredding efficiency (Fig. A7). While high male fitness typically led to suppression of both populations, female fitness had little impact (Fig. A8). Furthermore, higher levels of spillover and high shredding efficiency, led to a wider range of parameters where suppression occurred in both populations.

### Combinations of M-linkage and shredding can suppress a target population without significant impact on a neighboring population

The results presented so far (Table 1, Figs. 2-4) suggest that the m-shredders (E) can be highly efficient at population suppression when they are relatively tightly linked to the M locus and when the shredding efficiency is high. However, there may exist scenarios when little or no impact of the released transgene on neighboring populations is preferred. We found that suppression of the target population, where the shredding allele was introduced, could be achieved without suppression of (or editor gene persistence within) a neighboring population (Figs. 5, A7, and A8). However, as expected for any self-limiting population suppression method, even when population 1 was driven to local elimination with repeated releases, if population 2 persisted, the continued spillover of population 2 back into population 1 led to the rebound of population 1. Thus, continued release in population 1 is required to prevent migrants from population 2 to expand in a “vacuum” environment. Spatially distinct populations may be buffered by a region such that reintroduction of population 2 is unlikely once population 1 has been suppressed, which will be explored in future work incorporating space explicitly. Importantly, a range of linkage led to (temporary) local elimination of population 1, but without persistence of the editor gene in population 2 (Fig. 5). Low to intermediate male fitness was necessary for the effects of the SRD to be confined to a target population (Figs. A7 and A8). The ability to only drive the target population to local elimination but have only temporary impact on a neighboring population was favored in scenarios with combinations of the following: high shredding efficiency, low to intermediate male fitness, and high release fraction. To further assess the possibility of generating effective M-linked m-shredders, for fixed combinations of shredding efficiency and male fitness, we determine the minimum and maximum linkage required to drive population 1 below a low threshold (5%) with 30 repeated releases with release fraction of 0.5*×* (Fig. A9) or 2*×* (Fig. 6) while the editor gene does not persist in the neighboring population. These results indicate that it is possible to modulate the invasiveness of the SRD by manipulating M-linkage. Overall, these findings were consistent with parameter sensitivities observed for the single population model (Fig. A10).

**Figure 5.**
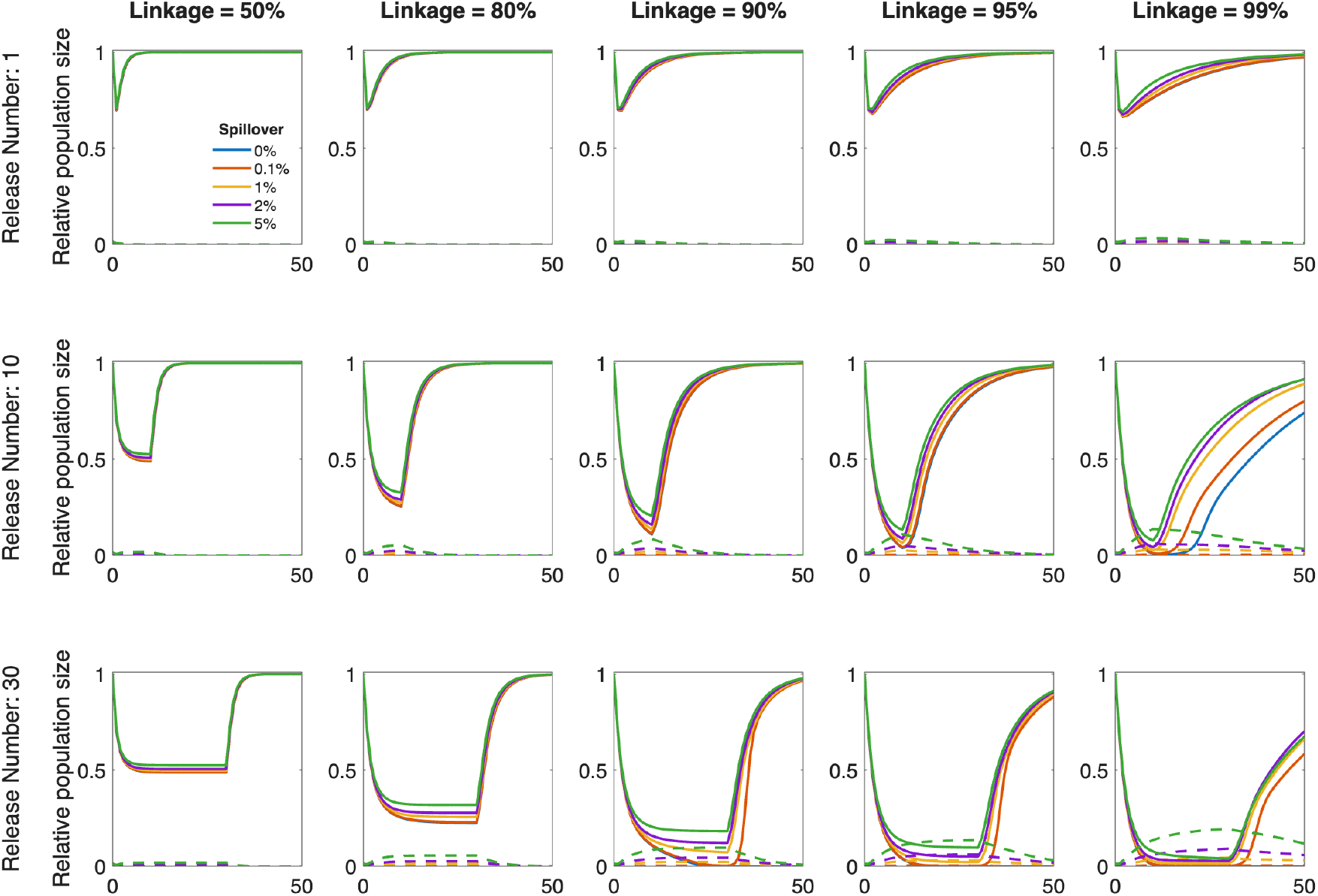
Dynamics of mosquito populations following a single release (top row) and repeated release (second row with 10 releases; and bottom row with 30 releases) of M^E^/m^a^ males under varying linkage and spillover. Relative population size (solid lines = adult female mosquitoes relative to wildtype population at equilibrium in the target population, population 1; dashed lines = total adult male and female mosquitoes carrying at least one copy of the editor gene relative to the current total adult mosquito population in neighboring population, population 2) following the introduction of M^E^/m^a^ males released at 0.5*×* the equilibrium male population. Linkage varies across panels from left to right: 50% (or autosomal), 80%, 90%, 95%, and 99% (nearly fully linked). Line color denotes spillover per generation (blue = 0%, red 0.1%, orange 1%, purple 2%, and green 5%). Shredding efficiency is 95%. Fitness of male genotypes with shredding allele is 50% that of wildtype genotype. Relative population size in population 1 for 0% spillover (blue lines) is nearly identical to that for 0.1% spillover under many conditions, so the lines may not be visually apparent. In the absence of spillover, no editor gene leaks into population 2, so the dashed blue lines are always zero. Unless otherwise specified, baseline parameter values from Table A1 are used.

**Figure 6.**
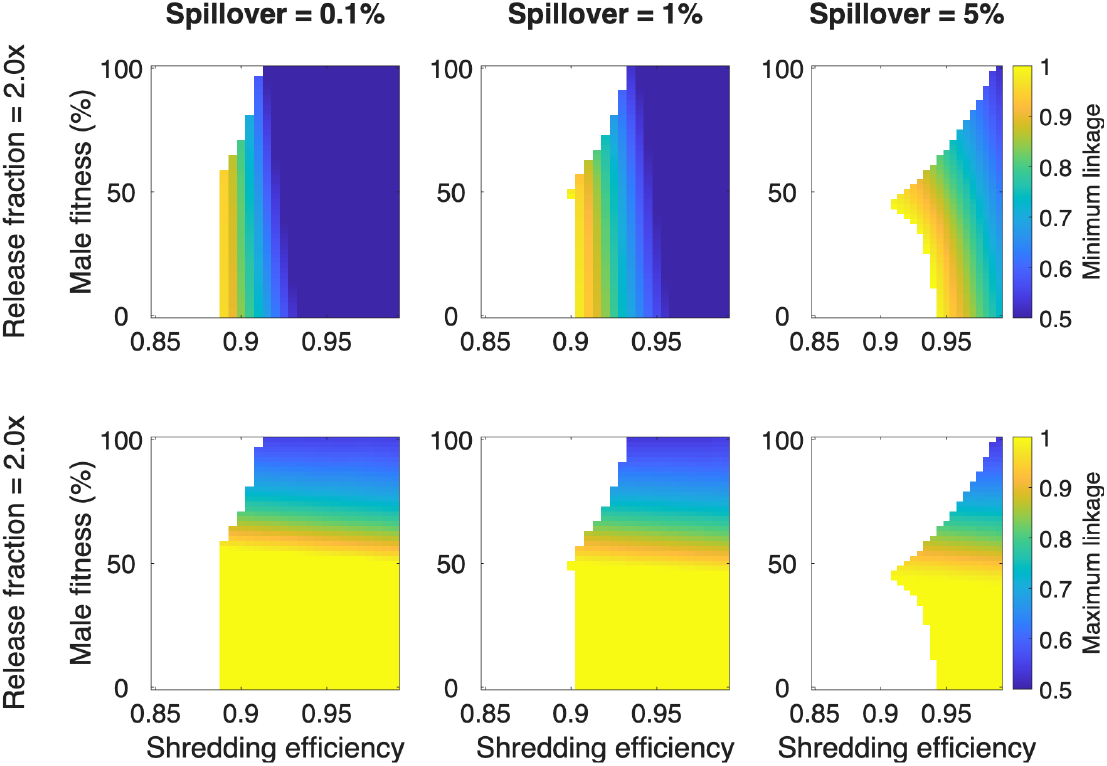
Minimum (top row) and maximum (bottom row) linkage for suppression of population 1 while maintaining persistence of population 2 without the presence of E alleles following 30 repeated releases of of M^E^/m^a^ males with varying shredding efficiency and male fitness. Release fraction of 0.5*×* male wildtype equilibrium under (*left to right* ) 0.1%, 1%, and 5% spillover. Unless otherwise specified, baseline parameter values from Table A1 are used.

### Population suppression is sensitive to linkage, shredding efficiency, and population growth rate

The required release fraction for population suppression (measured as reduction by 95% of the wildtype equilibrium population) was highly sensitive to parameters that dictated the level of genotypes in the following generation such as shredding efficiency and linkage (Fig. A10). For each population growth rate, a minimum shredding efficiency was required, regardless of release fraction or frequency, for suppression to be possible at all. For shredding efficiency above this level, even relatively low release fractions were often sufficient to reach population suppression, especially given higher linkage. With a larger population growth rate, the ability for strong population suppression was diminished because the higher number of offspring from successfully mated females allowed for persistence above the threshold of 95% reduction (Fig. 7).

**Figure 7.**
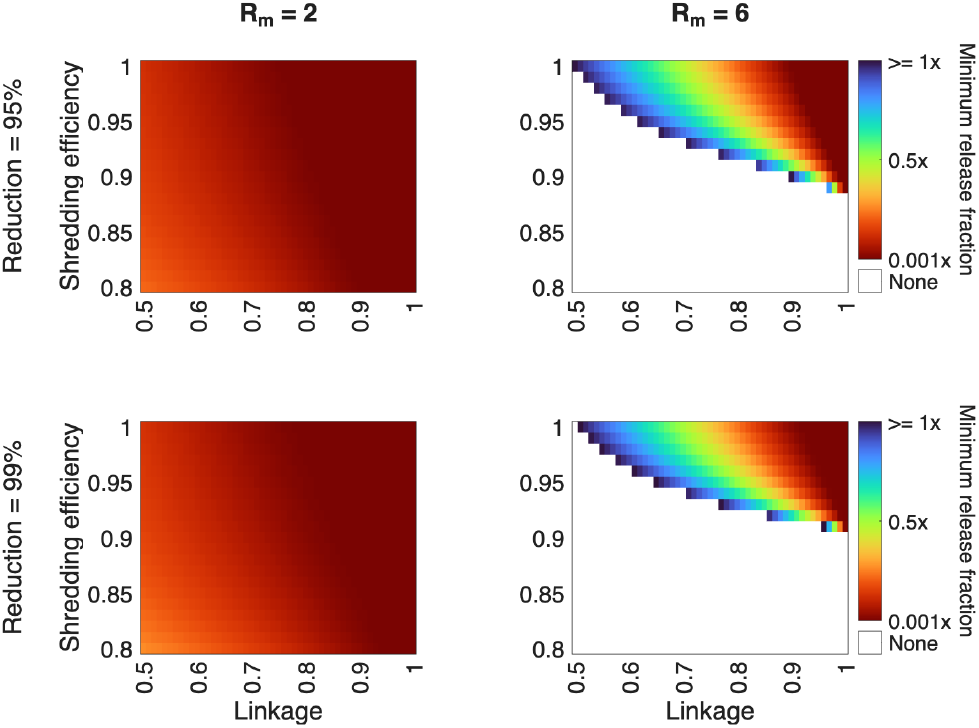
Minimum release fraction to achieve 95% (top row) or 99% (bottom row) reduction as shredding efficiency and linkage are varied with low intrinsic growth rate (left column, *R*_*m*_ = 2) or standard intrinsic growth rate (right column, *R*_*m*_ = 6). White indicates that no release fraction tested (from 0.001*×* to 10*×*) was sufficient. Black indicates that release fraction between 1 and 10*×* was required. See Section A.3 for details on intrinsic growth rate calculation. Unless otherwise specified, baseline parameter values from Table A1 are used.

## Discussion

Sex ratio distortion, as shown by the synthetic X-shredders developed for *Anopheles gambiae* [5, 6] are significantly more efficient at population suppression than the classic sterile insect technique (SIT) and incompatibility insect technique (IIT) methods due to their multi-generational effect at reducing the female population. However, Y-linked X-shredders have not been successfully engineered due to meiotic sex chromosome inactivation that renders them inactive during spermatogenesis [13, 14]. Meiotic sex chromosome inactivation is not expected in *Aedes aegypti* and other culicine mosquitoes as they contain homomorphic sex chromosomes [27, 31]. Here we developed deterministic and stochastic models to examine M-linked sex ratio distorters (SRDs, or m-shredders) by incorporating new features stemming from the homomorphic nature of the Aedes sex chromosomes and explored a broad range of parameter combinations that impact the efficiency and local confinement of these M-linked m-shredders.

As mentioned earlier, the idea of a Y-linked X-shredder or other Y-linked SRDs have been examined through models previously; however, all the models assume autosomal or full Y-linkage [8, 9, 10, 11]. While more simplistic in formulation than the models of Deredec *et al*. [11] and Beaghton *et al*. [9], our model allows for variable M-linkage with the m-shredder such that the impact on target populations as well as neighboring populations is tunable. As the M-linkage is determined by the location of the m-shredder relative to the M locus [21], this parameter can be readily manipulated in strain development.

According to our deterministic model, the time to suppression and final population size are very similar for shredders with 98-100% M-linkage, achieving rapid population suppression with a single release of 10% release fraction when the shredding efficiency is *≥* 95%. Moreover, population suppression is possible for a wide range of M-linkage and shredding efficiency, and the inclusion of demographic stochasticity further enhances the likelihood of population suppression as demonstrated by the stochastic simulations. These results suggest that it is critical to develop effective m-shredding editors to achieve population suppression, and it is not necessary to insert the m-shredder into the challenging repeat-rich M-locus to achieve highly efficient population suppression as 98% M-linkage can be accomplished by inserting the m-shredders in a *>*100 Mb genomic region [21]. For biologically obtainable values of shredding efficiency and linkage, the necessary release fraction to reduce the population by 95% is much below comparable population suppression techniques (e.g., those found in [10]) such as Y-Linked Editor, Release of Insects carrying Dominant autosomal Lethal genes (RIDL), or autosomal X-shredder (Table 1).

As it is possible to modulate the effectiveness of the shredder by changing the M-linkage, we further explored the parameter space that enables confined population suppression. We identified a range of M-linkage (dependent on given shredding efficiency) that can suppress a target population without significant impact on a neighboring population. However, continued release in the target population, perhaps in a spatial buffer zone, is required to prevent migrants from the neighboring population to expand in a “vacuum” environment.

In summary, we established mathematical models and evaluated in silico M-linked m-shredders with a range of efficiencies and confinement properties, which expands the toolbox that could help accommodate the varied needs facing global mosquito-borne infectious disease control programs. Our model and analysis provides an in-depth understanding of which parameters are most important for developing successful M-linked driving constructs aimed at population suppression. Furthermore, the framework developed here is suitable for any species with homomorphic sex chromosomes.

## Supporting information

Supplemental Methods and Results

