## Supplemental Methods and Results for "Modeling population control via tunable sex ratio distorter gene drives in *Aedes aegypti*, an arboviral vector with homomorphic sex chromosomes"

#### A Supplementary Methods

Our population-level mathematical model for the dynamics of populations of genetically distinct *Aedes* mosquito populations is represented in Figure A1 and tracks the number of juveniles and adults of each genotype. Each genotype is represented by two loci: one representing sex determination (M or m) and the other representing an 'editor' gene (E or a, where a is wildtype). In each generation, the juvenile population increases through birth, where the number and genotype of offspring is determined by the genotypes and abundance of the adult mating pairs (represented through the birth function $B_i(M_j, F_i)$, described in Section A.3). In a single generation, all present juveniles either develop into adults or die. The adult population increases due to the development of juveniles, which incorporates both density-dependent (via $\alpha$) and density-independent (via $\theta$) mortality as well as fitness of the each genotype, $f_i$. The adult population decreases from natural mortality, $\mu$. In contrast to juveniles, some fraction of the adult population can persist between generations, generating the potential for overlapping generations. In addition to the mortality and fitness parameters, the birth function incorporates parameters for: linkage, m-shredding efficiency, dominance coefficients, selection coefficients, and fertilized eggs per wildtype female per generation. See Table A1 for parameter descriptions and values.

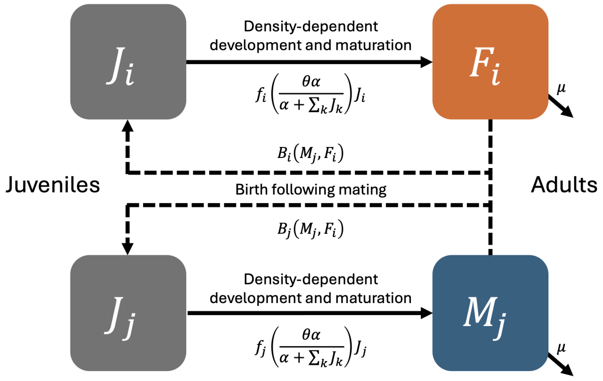


Figure A1: Schematic of conceptual model which is implemented deterministically and stochastically. Each genotype (represented by the subscript) has a corresponding juvenile ($J_i$ or $J_j$) and adult ($F_i$ or $M_j$) population. Details of the deterministic and stochastic representations are found in Section A.1.

##### A.1 Mathematical representations

**Mean-field deterministic representation.** We express our compartmental model via a mean-field approximation in two ways: ordinary differential equations (ODEs) and difference equations.

The ODEs for each juvenile genotype $k$, female genotype $i$, and male genotype $j$ are given by

$$\begin{aligned}
\overbrace{\frac{dJ_k}{dt}}^{\text{change in juvenile population}} &= \overbrace{B_k(M_j, F_i \;\forall\; j, i)}^{\text{birth function of juveniles}} - \overbrace{J_k}^{\text{juvenile loss}}, \\
\underbrace{\frac{dF_i}{dt}}_{\text{change in adult female population}} &= \underbrace{f_i\left(\frac{\theta\alpha}{\alpha + \sum_k J_k}\right) J_i}_{\text{development of surviving female juveniles}} - \underbrace{\mu F_i}_{\text{adult female loss}}, \\
\underbrace{\frac{dM_j}{dt}}_{\text{change in adult male population}} &= \underbrace{f_j\left(\frac{\theta\alpha}{\alpha + \sum_k J_k}\right) J_j}_{\text{development of surviving male juveniles}} - \underbrace{\mu M_j}_{\text{adult male loss}}.
\end{aligned} \tag{1}$$

The difference equations are similarly given by

$$\begin{aligned}
J_k(t+1) &= B_k(M_j(t), F_i(t) \;\forall\; j, i), \\
F_i(t+1) &= F_i(t) + f_i\left(\frac{\theta\alpha}{\alpha + \sum_k J_k(t)}\right) J_i(t) - \mu F_i(t), \\
M_j(t+1) &= M_j(t) + f_j\left(\frac{\theta\alpha}{\alpha + \sum_k J_k(t)}\right) J_j(t) - \mu M_j(t).
\end{aligned} \tag{2}$$

**Stochastic representation.** As the deterministic representation constitutes a mean-field approximation wherein demographic fluctuations that become especially relevant for small population counts as well as temporal correlations whose importance too is amplified near extinction thresholds are neglected, we also utilize a stochastic representation of the compartmental model. Details of the stochastic implementation are found in Section A.5. Naturally, stochastic models are considerably more costly to run since obtaining decent statistics requires multiple repetitions and averaging over distinct temporal histories. Yet these repeated realizations also yield relevant probabilistic information such as estimated likelihoods of successfully suppression of undesired genotypes.

#### A.2 Intrinsic population growth.

For the wildtype population, our system collapses down to two genotypes ($\mathrm{M^a/m^a}$ and $\mathrm{m^a/m^a}$) and thus only four equations (for juvenile males, $J_{\mathrm{M^a/m^a}}$; juvenile females, $J_{\mathrm{m^a/m^a}}$; adult males, $M_{\mathrm{M^a/m^a}}$; and adult females, $F_{\mathrm{m^a/m^a}}$). In this case, there are two equilibria, the extinction equilibria ($J_{\mathrm{M^a/m^a}} = 0$, $J_{\mathrm{m^a/m^a}} = 0$, $M_{\mathrm{M^a/m^a}} = 0$, $F_{\mathrm{m^a/m^a}} = 0$) and endemic equilibria given by

$$\left(J_{\mathrm{M^a/m^a}} = \frac{\alpha}{4\mu}(f_a\beta\theta - 2\mu),\; J_{\mathrm{m^a/m^a}} = \frac{\alpha}{4\mu}(f_a\beta\theta - 2\mu),\; M_{\mathrm{M^a/m^a}} = \alpha\left(\frac{f_a\theta}{2\mu} - \frac{1}{\beta}\right),\; F_{\mathrm{m^a/m^a}} = \alpha\left(\frac{f_a\theta}{2\mu} - \frac{1}{\beta}\right)\right).$$

The stability of each equilibrium is determined by the whether $R_m = \frac{f_a\beta\theta}{2\mu}$ is greater or less than one. When $R_m < 1$, the extinction equilibrium is stable and when $R_m > 1$ the endemic equilibrium is stable. The numerical value of $R_m$ also gives an indication of the intrinsic growth rate of the population. Thus, given our standard choice of parameters ($\beta = 12$, $f_a = 1$, $\theta = 1$, $\mu = 1$), this results in an intrinsic growth rate of $R_m = 6$, chosen for consistency with prior work [10].

#### A.3 Birth Function

The complexity of our model arises from inheritance of genetic alleles following mating and, thus, the generation of genetically diverse offspring from the birth function. As each parent produces haploid gametes, which combine to form diploid offspring upon mating, the combination of offspring from mating pairs may be more diverse than the individual parental genotypes alone. The probabilities of generating various offspring genotypes from any given parental combination are represented by a birth matrix, where two parameters determine the distribution of offspring genotype by mating pair: shredding efficiency, $s$, and linkage, $\ell$ (both discussed in detail below). The abundance of the offspring genotypes is determined by the abundance of parental genotypes in combination with the offspring probabilities from the individual mating pairs. The birth function for genotype $k$ is given by

$$B_k(M_j, F_i \;\forall\; j,i) = \frac{\beta}{\sum_k M_k} \sum_{ij} F_i M_j [\text{birth matrix}],$$

where $\beta$ is the average number of productive offspring per mating pair. See below for the birth matrix of all mating pairs. Furthermore, females mate only once during their lifespan, and we assume that males can mate, on average, with no more than five females. This constraint of the number of male matings ensures the opportunity for effective population reduction, particularly when the male population is small, i.e., a single male cannot allow population persistence by mating with tens or hundreds of females in a single generation.

**Birth matrix with shredding.** The mating pairs (left column) produce the distribution of offspring, which depends on the level of linkage ($\ell$) and the shredding efficiency ($s$, which determines $s_m$ and $s_f$) given by

| | $\mathrm{M^E/m^E}$ | $\mathrm{M^E/m^a}$ | $\mathrm{M^a/m^E}$ | $\mathrm{M^a/m^a}$ | $\mathrm{m^E/m^E}$ | $\mathrm{m^E/m^a}$ | $\mathrm{m^a/m^E}$ | $\mathrm{m^a/m^a}$ |
|---|---|---|---|---|---|---|---|---|
| $\mathrm{M^E/m^E + m^E/m^E}$ | $s_m$ | 0 | 0 | 0 | $s_f$ | 0 | 0 | 0 |
| $\mathrm{M^E/m^E + m^E/m^a}$ | $\frac{s_m}{2}$ | $\frac{s_m}{2}$ | 0 | 0 | $\frac{s_f}{2}$ | $\frac{s_f}{2}$ | 0 | 0 |
| $\mathrm{M^E/m^E + m^a/m^E}$ | $\frac{s_m}{2}$ | $\frac{s_m}{2}$ | 0 | 0 | $\frac{s_f}{2}$ | $\frac{s_f}{2}$ | 0 | 0 |
| $\mathrm{M^E/m^E + m^a/m^a}$ | 0 | $s_m$ | 0 | 0 | 0 | $s_f$ | 0 | 0 |
| $\mathrm{M^E/m^a + m^E/m^E}$ | $\ell\ s_m$ | 0 | $(1-\ell)\ s_m$ | 0 | $(1-\ell)\ s_f$ | 0 | $\ell\ s_f$ | 0 |
| $\mathrm{M^E/m^a + m^E/m^a}$ | $\frac{\ell\ s_m}{2}$ | $\frac{\ell\ s_m}{2}$ | $\frac{(1-\ell)\ s_m}{2}$ | $\frac{(1-\ell)\ s_m}{2}$ | $\frac{(1-\ell)\ s_f}{2}$ | $\frac{(1-\ell)\ s_f}{2}$ | $\frac{\ell\ s_f}{2}$ | $\frac{\ell\ s_f}{2}$ |
| $\mathrm{M^E/m^a + m^a/m^E}$ | $\frac{\ell\ s_m}{2}$ | $\frac{\ell\ s_m}{2}$ | $\frac{(1-\ell)\ s_m}{2}$ | $\frac{(1-\ell)\ s_m}{2}$ | $\frac{(1-\ell)\ s_f}{2}$ | $\frac{(1-\ell)\ s_f}{2}$ | $\frac{\ell\ s_f}{2}$ | $\frac{\ell\ s_f}{2}$ |
| $\mathrm{M^E/m^a + m^a/m^a}$ | 0 | $\ell\ s_m$ | 0 | $(1-\ell)\ s_m$ | 0 | $(1-\ell)\ s_f$ | 0 | $\ell\ s_f$ |
| $\mathrm{M^a/m^E + m^E/m^E}$ | $(1-\ell)\ s_m$ | 0 | $\ell\ s_m$ | 0 | $\ell\ s_f$ | 0 | $(1-\ell)\ s_f$ | 0 |
| $\mathrm{M^a/m^E + m^E/m^a}$ | $\frac{(1-\ell)\ s_m}{2}$ | $\frac{(1-\ell)\ s_m}{2}$ | $\frac{\ell\ s_m}{2}$ | $\frac{\ell\ s_m}{2}$ | $\frac{\ell\ s_f}{2}$ | $\frac{\ell\ s_f}{2}$ | $\frac{(1-\ell)\ s_f}{2}$ | $\frac{(1-\ell)\ s_f}{2}$ |
| $\mathrm{M^a/m^E + m^a/m^E}$ | $\frac{(1-\ell)\ s_m}{2}$ | $\frac{(1-\ell)\ s_m}{2}$ | $\frac{\ell\ s_m}{2}$ | $\frac{\ell\ s_m}{2}$ | $\frac{\ell\ s_f}{2}$ | $\frac{\ell\ s_f}{2}$ | $\frac{(1-\ell)\ s_f}{2}$ | $\frac{(1-\ell)\ s_f}{2}$ |
| $\mathrm{M^a/m^E + m^a/m^a}$ | 0 | $(1-\ell)\ s_m$ | 0 | $\ell\ s_m$ | 0 | $\ell\ s_f$ | 0 | $(1-\ell)\ s_f$ |
| $\mathrm{M^a/m^a + m^E/m^E}$ | 0 | 0 | $\frac{1}{2}$ | 0 | 0 | 0 | $\frac{1}{2}$ | 0 |
| $\mathrm{M^a/m^a + m^E/m^a}$ | 0 | 0 | $\frac{1}{4}$ | $\frac{1}{4}$ | 0 | 0 | $\frac{1}{4}$ | $\frac{1}{4}$ |
| $\mathrm{M^a/m^a + m^a/m^E}$ | 0 | 0 | $\frac{1}{4}$ | $\frac{1}{4}$ | 0 | 0 | $\frac{1}{4}$ | $\frac{1}{4}$ |
| $\mathrm{M^a/m^a + m^a/m^a}$ | 0 | 0 | 0 | $\frac{1}{2}$ | 0 | 0 | 0 | $\frac{1}{2}$ |

In the case of shredding, the total gamete number remains the same ($s_m + s_f = 1$), such that the probabilities depend on $s$ as follows: $s_m = \frac{1}{2-s}$, $s_f = 1 - s_m = \frac{1-s}{2-s}$. Details discussed below in Section A.4 **Shredding**.

#### A.4 Parameterization

**Linkage.** Linkage ($\ell$) between the male-determining locus and the editor gene locus, which we allow to be variable, entails that offspring with the same combination of the male-determining and editor gene loci as in the paternal genes are more likely, since the loci tend to be inherited together.

Table A1: Description of key parameters and their standard values. Full range indicates the parameter was varied across the full range of biologically possible values.
♯Fitness varies by genotype and is determined by the selection and dominance coefficient associated with the genotype. Baseline values listed are used for both females and males.

| Symbol | Description | Baseline value | Range | Reference |
|---|---|---|---|---|
| $\beta$ | average number of offspring per mating | 12 | 4, 12, 24 | [10] |
| $\alpha$ | intensity of density-dependent mortality | 200 | - | [10] |
| $\theta$ | density-independent probability of juvenile survival | 1 | - | [10] |
| $\mu$ | natural mortality of adult mosquitoes | 1 | - | assumed |
| $\ell$ | linkage | 0.98 (98%) | [0.5,1] | full range |
| $s$ | shredding efficiency | 0.90 (90%) | [0,1] | full range |
| $f_a$ | maximum fitness of wildtype genotypes♯ | 1 | - | assumed |
| $f_E$ | maximum fitness of editor genotypes♯ | 0.9 | [0,1] | full range |
| $s_a$ | selection coefficient in wildtype genotypes | 0 | - | assumed |
| $s_E$ | selection coefficient in editor-carrying genotypes | 0.1 | [0,1] | full range |
| $h_{Mm}$ | dominance coefficient in males | 1 | - | [7] |
| $h_{mm}$ | dominance coefficient in females | 1 | - | [7] |
| $\sigma$ | spillover between populations | 0 (0%) | [0,0.05] | assumed |

When $\ell = 0.5$, there is neutral inheritance, as this assumes that E is autosomal and thus not linked to the sex locus. When $\ell > 0.5$, the loci are considered linked and offspring with gene combinations that are linked in those in the parental genomes are more probable. For $\ell = 1$, the linked genes are always inherited together.

**Shredding.** The shredding efficiency parameter ($s$) determines the extent of male bias. The total number of gametes remains the same but the lost female gametes are replaced by males. The parameters $s_m$ and $s_f$ in the birth matrix capture the fraction of offspring generated that are either male ($s_m$) or female ($s_f$) in a single mating process. Note that when there is no shredding of female offspring ($s = 0$), then $s_m = s_f = 0.5$, and hence there is no bias towards male offspring, while for any $s > 0$, then $s_m > \frac{1}{2} > s_f$, which results in more males than females in the next generation. As the total gamete number remains the same ($s_m + s_f = 1$), the fractions (in this case, probabilities) depend on $s$ as follows:

$$s_m = \frac{1}{2-s} \ , \qquad s_f = 1 - s_m = \frac{1-s}{2-s} \ .$$

**Release fraction and type.** To understand the ability of an introduced editor gene to elicit population suppression, we release adult male mosquitoes with the editor gene linked to the male-determining locus (we focus on $\mathrm{M^E/m^a}$ release as it is biologically more realistic when E is linked to M) and track how the population size is altered after 30 generations. We consider release fraction (0.1×, 0.5×, 1×, 2×, 5×) relative to the total adult male population at wildtype equilibrium ($\mathrm{M^a/m^a}$ and $\mathrm{m^a/m^a}$), which causes a temporary increase in the size of the adult population. We primarily consider a single release, but also consider multiple releases (one per generation) of the same fraction (fixed relative to the wildtype equilibrium) up to thirty generations.

#### A.5 Stochastic implementation

We construct an individual-based stochastic representation extending the mean-field deterministic model by tracking each juvenile and adult through generation time steps subject to the following three random processes with associated prescribed probabilities:

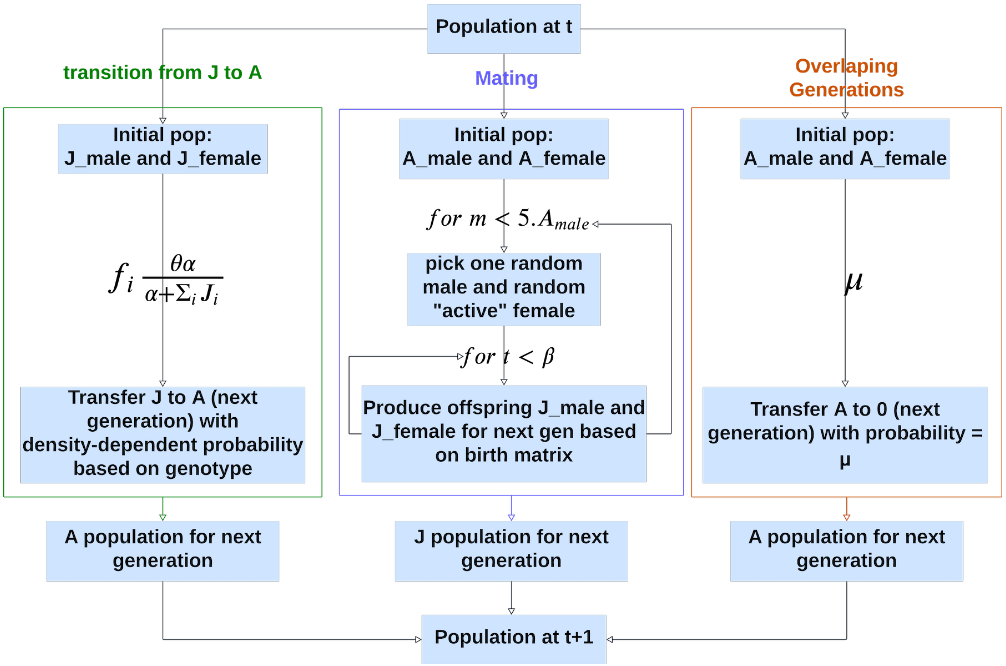


Figure A2: Flowchart of the dynamics of stochastic implementation in a single generation.

1. **Transition out of juvenile stage:** Juveniles of genotype $i$ become adults with the density-dependent probability:
$$f_i\left(\frac{\theta\alpha}{\alpha+\sum_i J_i}\right).$$
All other juveniles are lost.

2. **Production of offspring via mating:** We randomly select one male and one female from the population, with the restriction that each female may participate in mating only once per generation. Based on the genotypes of the chosen male and female, there are eight different probabilities which represent the chances of producing the distinct genotypes from a particular mating pair (as determined in the columns of the birth matrix). Each offspring is generated based on a uniform random draw on these prescribed probabilities. This process is iterated $\beta$ times. The resulting offspring may feature different genotypes. Next, we randomly select another male and female and repeat this mating process five times the total adult male population number, following the assumption of the maximum mating number for male individuals.

3. **Overlapping generations:** Adult individuals are removed from the population with the specified death probability, $\mu$, which is the same for all genotypes.

These steps are detailed in the stochastic simulation flow chart found in Fig. A2. We note that we describe each mathematical generation while a biological generation (e.g., juvenile to juvenile) corresponds to two mathematical generations. This is an important detail since in the stochastic model with non-overlapping generations, genotypes released as adults will not appear in the subsequent adult mathematical generation. Unless otherwise specified, when we use generation, we refer to biological generation.

##### A.6 Choice of parameterization for stochastic implementation

To ensure consistency between the parameterization of the deterministic and stochastic implementations of the model, we utilize the birth matrix with no bias towards male offspring ($s = 0$) and neutral linkage ($\ell = 1/2$), narrow the region of our parameter space by fixing the parameters $\theta = 1$, $\alpha = 200$, and $\mu = 1$ (chosen for consistency with previous work [10]), and vary the number of produced offspring $\beta$ (the number of fertilized eggs per mating per generation) in the range $6 \ldots 12$ in both mean-field and stochastic simulations. For each set of parameters examined, for comparison to the stochastic model, we numerically solve the deterministic ODE model using Runge-Kutta and average over 200 independent Monte Carlo simulations runs for the stochastic model. Figure A3 shows the dynamics of the total adult population over 60 generations subject to different values of $\beta$ for both the stochastic and mean-field deterministic simulations. By setting the initial population to 200 individuals, equally distributed across all genotypes, in both models, we find that both models reach their quasi-stationary population levels after roughly 10 generations. The ensuing quasi-stationary populations are the same in the mean-field and stochastic simulations.

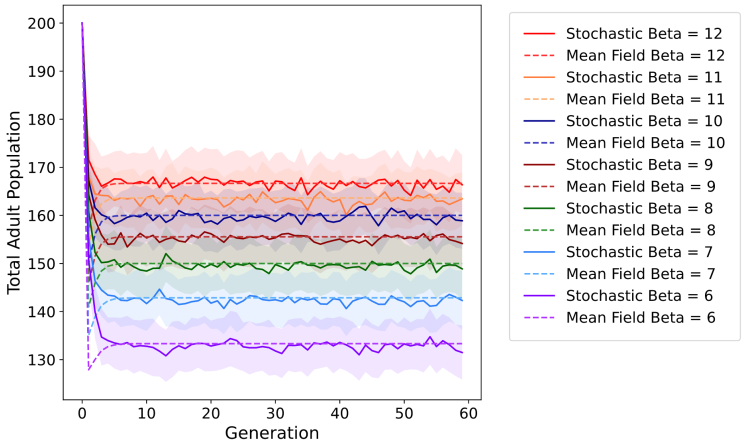


Figure A3: Adult population dynamics from the stochastic (solid line) and mean-field (dashed) model simulations across 60 generations for varying $\beta$ values. Parameters: $\theta = 1$, $\mu = 1$, $\alpha = 200$, $\ell = 0.5$ and $s = 0$. The initial population size is set to 200 individuals, evenly split between adult populations of each genotype. Stochastic simulations are averaged over 200 independent trials.

Figure A4 shows the (quasi-)stationary population levels after 60 generations as a function of the number of fertilized eggs per mating per generation ($\beta$). The error bars for $n = 200$ trials of the stochastic model represent the standard deviation of the stationary population data. This demonstrates that across different number of fertilized eggs the resulting stationary populations are consistent between the deterministic and stochastic models.

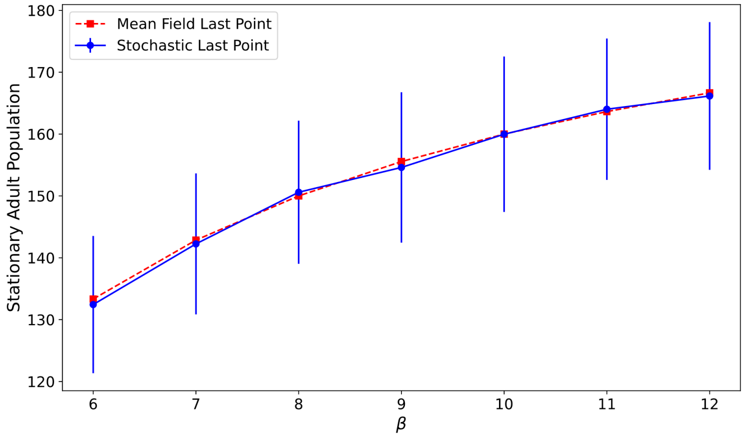


Figure A4: Total adult population after 60 generations for different $\beta$ resulting from integrating the mean-field rate equations (1) and $n = 200$ independent Monte Carlo simulation runs for the corresponding stochastic model. Parameters used: $\theta = 1$, $\mu = 1$, $\alpha = 200$, $\ell = 0.5$ and $s = 0$. The initial population size is set to 200 individuals, evenly split between adult populations of each genotype. Stochastic results are averaged over $n = 200$ independent trials. Error bars represent the standard deviation.

### B Supplemental Results

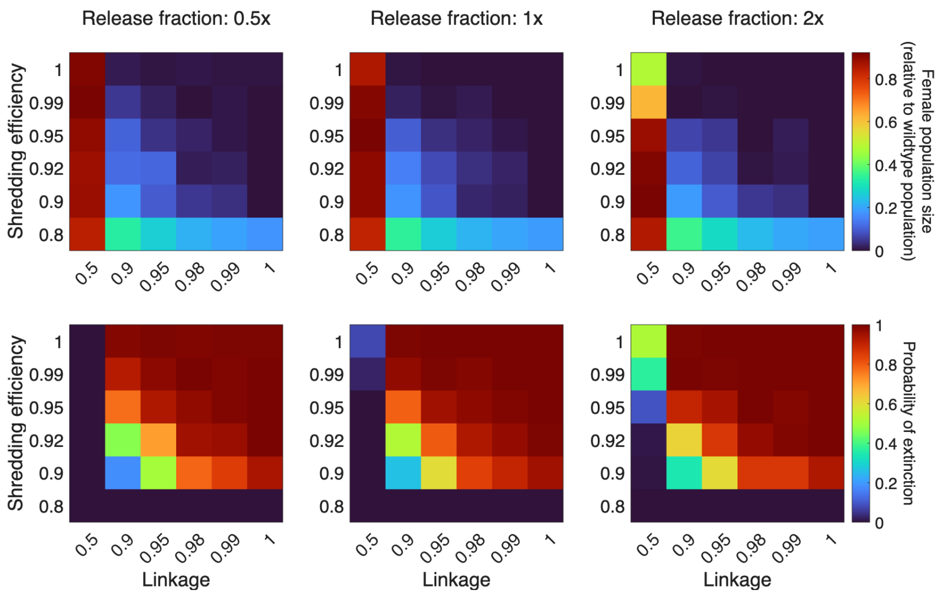


Figure A5: Female population size relative to wildtype equilibrium at 30 generations (top panels) and probability of extinction within 30 generations (bottom panels) with five releases of $M^E/m^a$ males for varying linkage and shredding efficiency as obtained from our stochastic representation. From left to right, the release fraction increases: 0.5×, 1×, 2×. Note the non-linear scaling of the axes. Fitness of genotypes with shredding allele is 100% that of wildtype genotype as it was determined that fitness costs of genotypes containing the shredding allele had minimal effect on the results for the deterministic representation for 80% to 100% fitness (Fig. A6). Stochastic output averaged over 200 replicate simulations with release starting after the wildtype population reached equilibrium. Unless otherwise specified, baseline parameter values from Table A1 are used.

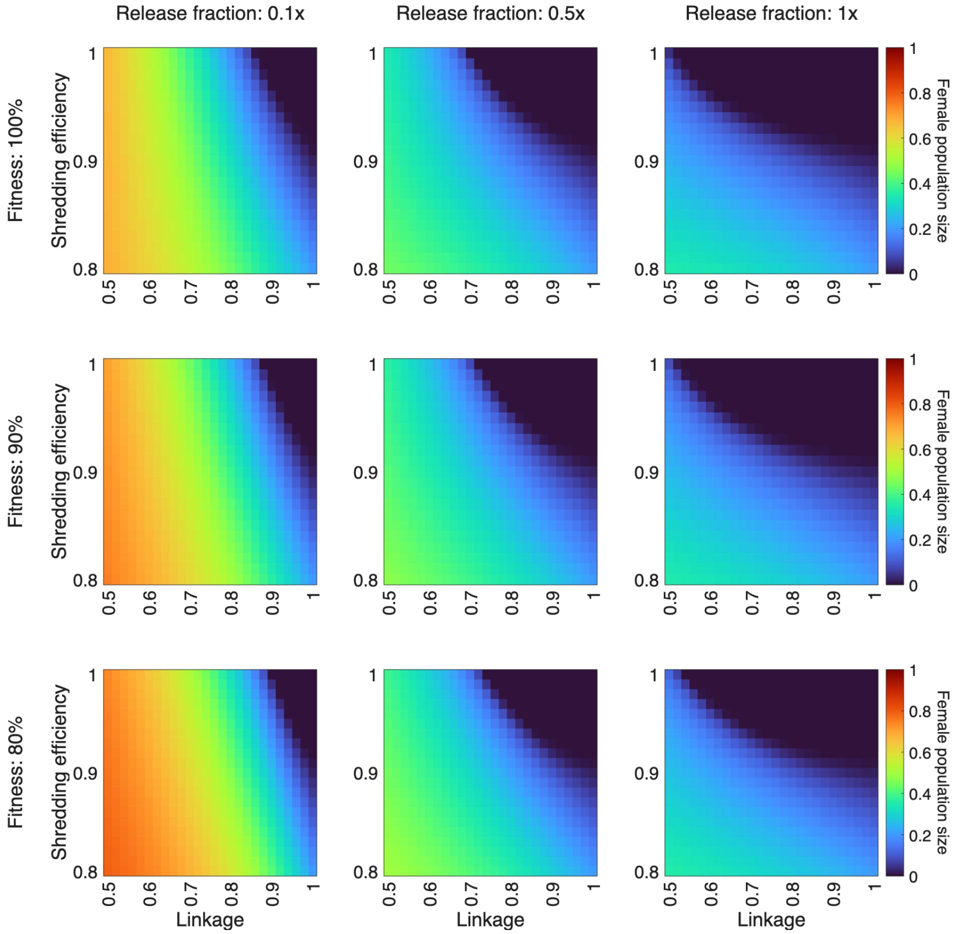


Figure A6: Female population size (relative to wildtype equilibrium) at 30 generations with 30 repeated releases of $M^E/m^a$ males for varying linkage and shredding efficiency. From left to right, the release fraction increases: 0.1×, 0.5×, 1×. From top to bottom, fitness of genotypes with shredding allele decreases: 100%, 90%, 80%. Unless otherwise specified, baseline parameter values from Table A1 are used.

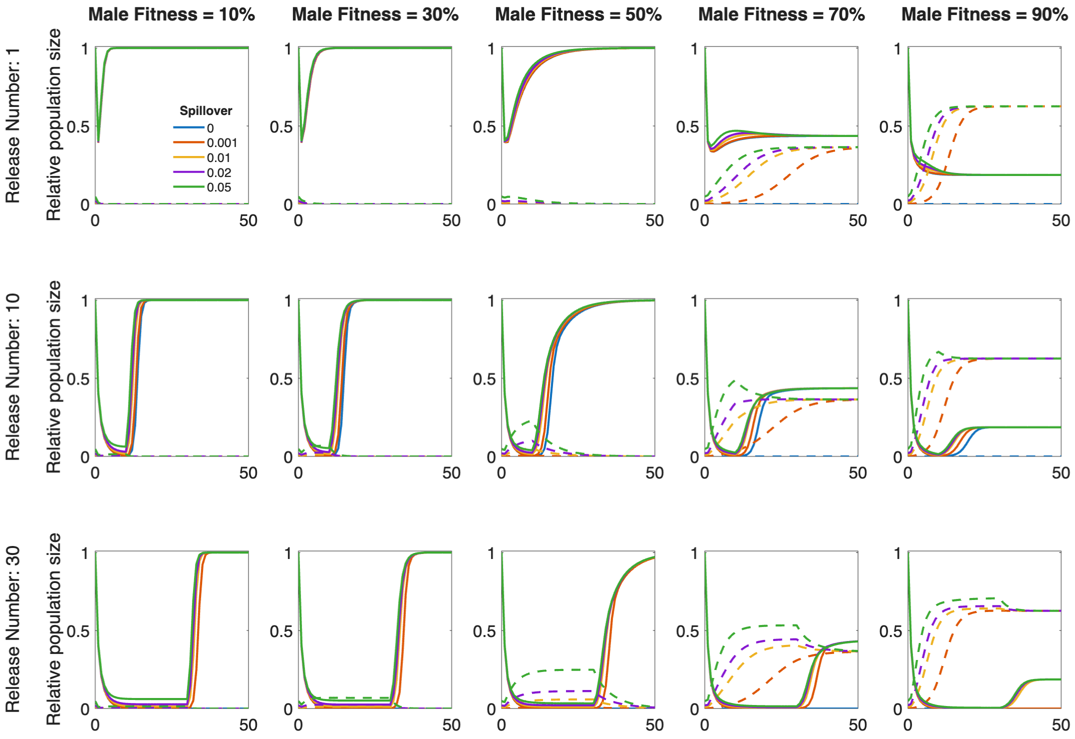


Figure A7: Dynamics of mosquito populations following a single release (top row) and repeated release (second row with 10 releases; and bottom row with 30 releases) of $M^E/m^a$ males under varying linkage and spillover. Relative population size (solid lines = adult female mosquitoes relative to wildtype population at equilibrium; dashed lines = total adult male and female mosquitoes carrying at least one copy of the editor gene relative to the current total adult mosquito population) following the introduction of $M^E/m^a$ males released at 2× the equilibrium male population. Fitness of male genotypes with shredding allele varies across panels from left to right: 10%, 30%, 50%, 70%, and 90%. Line color denotes spillover per generation (blue = 0%, red 0.1%, orange 1%, purple 2%, and green 5%). Shredding efficiency is 95%. Linkage is 90%. Unless otherwise specified, baseline parameter values from Table A1 are used.

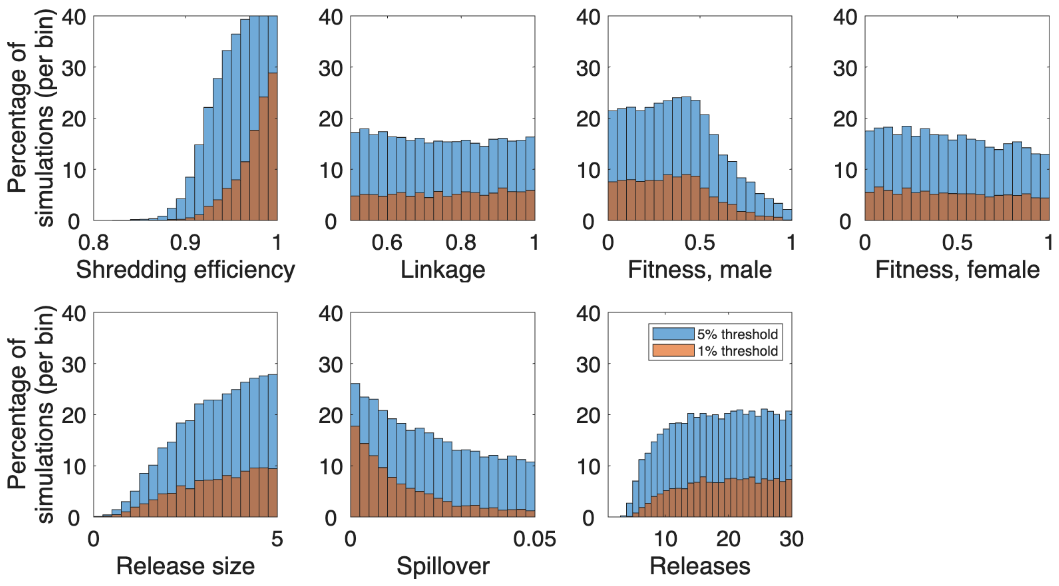


Figure A8: Values of shredding efficiency, linkage, fitness (male and female, uncorrelated), release fraction, spillover, and number of releases that lead to extinction in the target population (population 1) and persistence in the neighboring population (population 2) following release of $M^E/m^a$ males. Blue bars show for what parameters the size of population 1 drops by 95% while the size of population 2 remains within 5% of equilibrium, and red bars show for what parameters the size of population 1 drops by 99% while the size of population 2 remains within 1% of equilibrium. Unless otherwise specified, baseline parameter values from Table A1 are used.

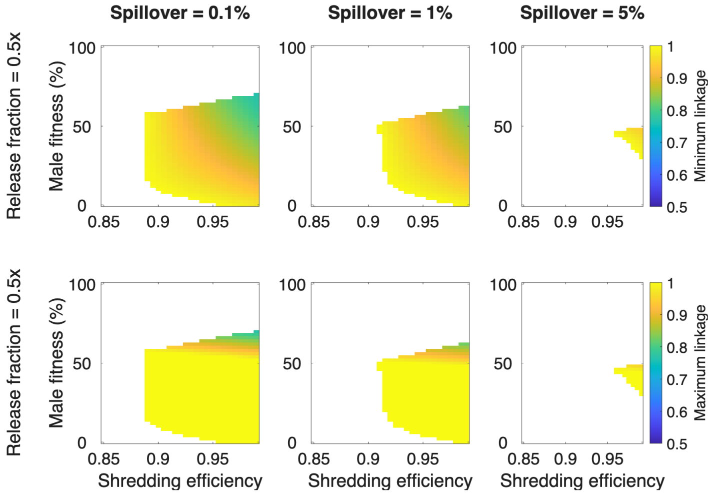


Figure A9: Minimum (top row) and maximum (bottom row) linkage for suppression of target (population 1) while maintaining persistence of neighboring population (population 2) without the presence of E alleles following 30 repeated releases of $M^E/m^a$ males with varying shredding efficiency and male fitness. Release fraction of 0.5× male wildtype equilibrium under (*left to right*) 0.1%, 1%, and 5% spillover. Unless otherwise specified, baseline parameter values from Table A1 are used.

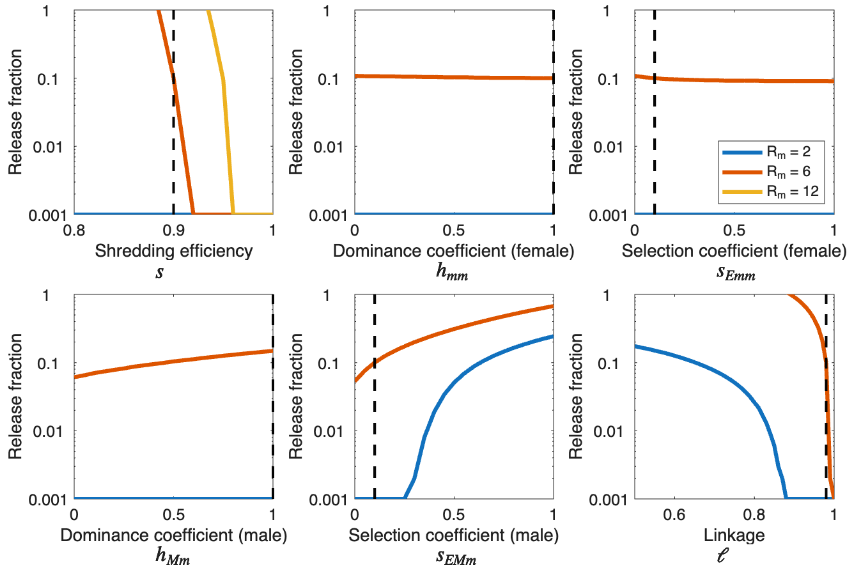


Figure A10: Univariate sensitivity analysis of key parameters. The minimum release fraction needed to achieve 95% reduction of the population after 30 generations of repeated release as shredding efficiency ($s$), dominance coefficient (female) ($h_{mm}$), selection coefficient (female) ($s_{Emm}$), dominance coefficient (male) ($h_{Mm}$), selection coefficient (male) ($s_{EMm}$), and linkage ($\ell$) are varied across the range specified in Table A1. Solid blue, red, yellow lines indicate intrinsic growth rates of $R_m = 2$, $R_m = 6$, and $R_m = 12$, respectively. Dashed black lines indicate baseline parameter value as listed in Table A1. All parameters except the one varied are held at baseline parameter values from Table A1 are used.
